# Synergistic amplifier reprograms biosensor behavior to facilitate detection of dioxin-like compounds

**DOI:** 10.1101/2024.06.26.600751

**Authors:** Chao Du, Yun Wang, Jiaheng Wang, Shunyi Li, Yuling Zhou, Li Yi, Jun Ge, Guimin Zhang

## Abstract

Biosensors are prominent detection tools for evaluating various analytical markers. However, these devices that meet practical application scenarios are still limited, and their potential remains to be developed. In these aspects, we attempted to design and construct a synergistic amplifier consisting of two components, namely, a transcriptional activation component mediated by the Gal3^C^ regulator and a transcription recruitment component exploiting the function of dCas9-(SpyC)_n_, which work together to enhance yeast biosensor performance. This synergistic amplifier increased the signal output by 5.1-folds and the detection limit by 4.4-folds, respectively, while maintaining adjustable sensitivity and avoiding noise introduction during signal amplification. We demonstrated the use of synergistic amplifier in detecting dioxin analogues, aiming to unlock the potential of predictive modulation, and found that it achieved progress in dynamic output and detection limit. This work show that synergistic amplifier is a new paradigm for expanding signal processing pathways within synthetic genetic networks, enabling cell-based biosensors for environmental monitoring.

## Introduction

Redesigning the molecular basis of life to achieve new functions has made whole-cell biosensors as research focus in the emerging field of synthetic biology (1–3). These genetic reprogramming detection devices have been extensively studied for customized signal processing within cells for various beneficial applications, such as clinical diagnosis (4,5), environmental and agricultural monitoring (6–8), and high-value goods, or industrial chemical production (9,10), etc. However, despite these promising technologies, they still face some challenges, such as sensing capabilities that are suitable for laboratory procedures but are often not suitable for practical applications due to lack of versatility, functional tunability, or stringent industrial conditions (11–13). To effectively manipulate the behavior of biosensors, strategies such as genome mining (14), evolution (15), logic gate (16), transcriptional or translational level regulation (11) have promoted closer integration of biosensors with practical applications. Although in some cases established methods often focus only on specific parameters of the dose-response curve (such as dynamical range, basal leakage, or response threshold), multidimensional parameter tuning can improve biosensor behavior more comprehensively. Therefore, there is a strong demand to develop new methods to improve biosensor architecture using multidimensional tuning schemes.

Similar to the role of signal amplifier building blocks in electronic circuits, signal amplifier building blocks have been widely used to reshape signal carriers in the form of voltage or current waveforms in electrical and electronic systems (17,18). Genetic signal amplifiers can reprogram such signal processing pathways within biosensor architectures, thereby facilitating predictive modulation of signal outputs. Over the past decade, the use of cascaded transcriptional activation and transcription recruitment processes have enabled the amplification of signals within synthetic gene networks, amenable to mathematical modelling and quantitative analysis. For example, Wan et al. described a modular, cascaded signal amplification method by designing multilayered transcriptional amplifiers and observed a 5,000-fold and 750-fold increase in detection limit and output, respectively (11). Fan et al. recently combined the GPCR signaling pathway with the galactose regulatory system and incorporated a positive feedback loop into an engineered yeast biosensor, achieving a 100-fold improvement in LOD and a 41 times increase in maximum output level compared to the original sensor (19). Therefore, the cascade transcriptional activation effect is attributed to the stepwise enhancement of high-gain transcriptional processes and is therefore considered to be an important factor in signal amplification. Strikingly, as reported by Tanenbaum et al., the repetitive peptide array SunTag can recruit multiple copies of antibody fusion proteins, thereby enabling redesigned biological outputs (20). Subsequently, Zhai et al. developed a CRISPR-mediated protein-tagging signal processing system that can simultaneously activate and inhibit multiple genes in *Saccharomyces cerevisiae*; the gene activation and repression efficiencies are as high as 34.9-fold and 95%, respectively (21). This suggests that transcriptional recruitment promotes localization of transcriptional effectors with higher concentrations, which appear to be a key mechanism for signal amplification.

Among biosensing devices, whole-cell biosensors have been demonstrated to be feasible for detecting and quantifying dioxin-like compounds (DLCs), characterized by their bioavailability, persistence, and association with various diseases (22–24). However, several obstacles limit whole-cell biosensors for DLCs detection in practical settings, due mainly to their low dynamic output and unreliable sensitivity. Therefore, there is an urgent need to bring the performance of whole-cell biosensor closer to practical application. Inspired by cascade transcriptional activation and transcription recruitment, we developed a synergistic amplifier consisting of two components, a transcription activation component mediated by the Gal3^C^ regulator (a regulatory protein variant involved in the yeast *GAL* genetic network (25)) and a transcription recruitment component driven by dCas9-(SpyC)_n_ fusion (CRISPR/dCas9 technology (26) combined with SpyTag/SpyCatcher (abbreviated as SpyT/SpyC) technology (27)), which work together to exert a synergistic effect to enhance the performance of yeast biosensors. Considering the high degree of orthogonality and separability of the two components, each component was highly modular and a cumulative effect on signal amplification was observed. In the transcriptional activation component, synthetic promoters were engineered to specifically respond to the receptor protein complexes capable of sensing DLCs. The potency of the synthetic promoter and the transcriptional activation capacity of the regulator were also tunable. In the transcriptional recruitment component, the number of repeating SpyC units, the regulatory capacity of the transcriptional activator, and the target region of gRNA were all optimized to enhance sensor behavior. It was shown that the resulting engineered sensor embedded with a synergistic amplifier exhibits significant improvements in dynamic output, and maintains tunable sensitivity and detection limit. In addition, experiments involving the detection of dioxin analogues further demonstrated the signal amplification effect of the synergistic amplifier structure. These advances illustrated that the integration of a synergistic amplifier enables the programmable behavior of yeast biosensors, resulting in enhanced detection of persistent toxic substances.

## Materials and methods

### Materials

*E. coli* XL-gold or DH5α was stored in our laboratory and used for gene cloning. *Saccharomyces cerevisae* W303-1a was used as a host for analyzing genetic circuits, whereas EBY100 was used as an initial strain for constructing yeast biosensors. DNA polymerases, gel extraction kit and plasmid purification kit were purchased from Vazyme (Nanjing, China). T5 exonuclease, restriction endonuclease, and T4 DNA ligase were purchased from New England Biolabs (NEB, USA). *E. coli* was cultured using Luria–Bertani (LB) medium (0.5 % yeast extract, 1% tryptone and 1% NaCl) supplemented with 100 μg/mL ampicillin. Yeast extract peptone dextrose (YPD) medium (1% yeast extract, 2% peptone, 2% glucose), or synthetic complete (SC) medium (0.67% yeast nitrogen base, 2% glucose appropriate, 0.12% dropout supplements, pH 5.5) were used for the yeast cultivation. AhR agonist, β-naphthoflavone (β-NF, CAS Number 6015-87-2) was purchased from Macklin Biochemical Co. Ltd (Shanghai, China), 2,3,7,8-Tetrachlorodibenzo-*p*-dioxin (TCDD, CAS Number 1746-01-6) was purchased from Wellington Laboratories Inc. (Guelph Ontario, Canada), benzo[a]pyrene (BaP, CAS Number 50-32-8), 3,3’,4,4’,5-pentachlorobiphenyl (PCB-126, CAS Number 57465-28-8), and 3,3’,4,4’,5,5’-hexachlorobiphenyl (PCB-169, CAS Number 32774-16-6) were obtained from SCR-Biotech Co., Ltd (Shanghai, China). All chemicals and reagents are of analytical grade and commercially available.

The human *AhR* and *ARNT* genes were derived from the genome of *S. scerevisiae* BLYAhS (28). CRISPR/Cas9 system and yeast-enhanced green fluorescent protein (yEGFP) gene were kindly provided by Prof. Shuobo Shi from Beijing University of Chemical Technology. Based on the genome of *S. cerevisiae* W303-1a, regulatory protein genes involved in the yeast GAL genetic network, including *Gal4* (Gene ID: 855828), *Gal3* (Gene ID: 851572), and its three mutants, were amplified by PCR. The multi-copy gene of SpyCatcher (abbreviated as SpyC) was derived from our previous study (29). Primers, guide RNAs (gRNAs), oligonucleotide synthesis, and DNA sequencing were done by Tsingke Biotech Co., Ltd (Beijing, China).

### Design of a prototype yeast biosensor for the detection of dioxin-like compounds (DLCs)

Yeast episomal plasmids pESD (CEN/ARS *ori*, TRP1 selection marker), pBEVY (2μ *ori*, LEU2 or URA3 selection marker), pRS425 (2μ *ori*, TRP1 selection marker), and yeast integrative plasmid YIPlac204 (LEU2 selection marker) served as the parental templates for the construction of derivative plasmids. The plasmids used in this study are summarized in **Table S1**. A prototype yeast biosensor, which includes a sensor module and a report module, was constructed for the detection of dioxin-like compounds (DLCs) in strain EBY100. Human *AhR* and *ARNT* genes were cloned into pESD to generate pESD-AhR-ARNT. This construct is responsible for the expression of AhR and ARNT, and the subsequent formation of the DLCs-AHR-ARNT ternary complex in the nucleus upon DLCs binding. For the report module, yEGFP was placed under the control of the synthetic promoter DRE5-P_Core_, which contains a pentameric dioxin response element (5’-TNGCGTG-3’) and a TATA box, core elements, and transcription start site (TSS) within P_Core_, designed according to a previous report (30). This construct was then ligated into pBEVY to generate pBEVY-DRE5-P_Core_-yEGFP. The two plasmids were co-transformed into 100 μL of EBY100 competent cells by electroporation (2.0 kV, 5 ms) to obtain the biosensor. To adjust the maximum intracellular output level, various core elements within the synthetic promoters that directly drive yEGFP expression were replaced by polymerase chain reaction (PCR) amplification. **Table S3** provides a list of all engineered strains and sensors used in this study.

### Construct designs for signal amplification using transcriptional activation effect

To test the transcriptional activation ability, the plasmids pESD-P_TEF1_-Regulator containing constitutively expressed regulators (Gal4, Gal3, Gal3^F237Y^, Gal3^D368V^, and Gal3^S509P^) were constructed and co-transformed into EBY100 with the plasmid pBEVY-P_GAL1_-yEGFP respectively, to determine the fluorescence intensities of yEGFP under glucose-supplemented cultivation conditions. The regulator with the strongest activation ability was then used as an amplifier to manipulate the genetic signal within the biosensor and was renamed as Gal3^C^. To evaluate the effect of Gal3^C^ on the performance of the yeast biosensor, an amplification module based on Gal3^C^-mediated transcriptional activation was inserted between the sensor module and the report module. Given the limitations of the EBY100 auxotrophic selection markers, the plasmid pBEVY-Gal3^C^-P_Core_-DRE5-P_GAL1_-yEGFP was constructed and co-transformed with pESD-AhR-ARNT to obtain the engineered sensor.

### Construct of a signal amplification enabled by the synergistic amplifier

A genetic circuit consisting of the yeast integrative plasmid YIPlac204-P_TEF1_-Gal3^C^, the yeast episomal plasmid pRS425-dCas9-(SpyC)_n_-gRNA, and the yeast episomal plasmid pBEVY-TA-SpyT-P_GAL1-10_-yEGFP was used to determine the enhancement of yEGFP output by CRISPR-mediated transcriptional recruitment. dCas9 represents a Cas9 variant harboring two mutations (D10A, H840A). dCas9-(SpyC)_n_, where n = 1, 2, 3, or 4, were obtained by overlap extension PCR. The guide RNAs (gRNAs) targeting the template or non-template DNA strand of the P_GAL1-10_ region were designed using the E-CRISP website (http://www.e-crisp.org/E-CRISP/index.html). The gRNA spacer sequences are listed in **Table S2**. The gRNA expression cassette is consisted of the P_SNR52_ promoter, gRNAs spacer sequence (20 bp), gRNA scaffold (76 bp), and *SUP4* terminator (21). The fusion sequences of transcriptional activator (TA) and SpyT, TA-SpyT, including VPR-SpyT, VP64-SpyT, and Med2-SpyT, were synthesized by Tsingke Biotech Co., Ltd. The P_GAL1-10_ sequence was obtained by PCR reaction using the parental plasmid pESD as a template (31). pRS425-dCas9-(SpyC)_n_-gRNA and pBEVY-TA-SpyT-P_GAL1-10_-yEGFP were co-transformed into W3031a competent cells that pre-integrated with YIPlac204-P_TEF1_-Gal3^C^ to obtain the corresponding engineered strains (**Table S3**). Among the strains with the highest fluorescence intensity, the best performing genetic circuit, which consists of the synergistic action of the transcriptional activation element (mediated by the Gal3^C^ regulator) and the transcriptional recruitment component (driven by the dCas9-(SpyC)_n_ fusion), was identified, and the corresponding TA and gRNA spacer sequences were used in subsequent experiments. For ease of description, the term synergistic amplifier is used to represent the synergistic action between the transcriptional activation component and the transcription recruitment element.

To evaluate the effect of the synergistic amplifier on the performance of yeast biosensors. we engineered sensor constructs, specifically pESD-AhR-ARNT-dCas9-(SpyC)_n_-g3, pBEVY-SpyT-Med2-*IGG6*-Gal3^C^-P_Core3_-DRE5-P_GAL1_-yEGFP plasmids. The dCas9-(SpyC)_n_-g3 fragment (including the expression cassette of dCas9-(SpyC)_n_ and g3 guide RNA) was PCR amplified from pRS425-dCas9- (SpyC)_n_-gRNA and inserted into pESD-AhR-ARNT via homologous recombination. Likewise, the SpyT-Med2-*IGG6* fragment was PCR amplified from pBEVY-Med2-SpyT-P_GAL1-10_-yEGFP and inserted into pBEVY-Gal3^C^-P_Core3_-DRE5-P_GAL1_-yEGFP. *IGG6* is an intergenic sequence, as reported by Yue et al (32). Subsequently, various sensor constructs with synergistic amplifiers were obtained by co-transforming plasmids pESD-AhR-ARNT-dCas9-(SpyC)_n_-g3 and pBEVY-SpyT-Med2-*IGG6*-Gal3^C^-P_Core3_-DRE5-P_GAL1_-yEGFP into EBY100.

### Cell culture and data collection

Fluorescence analysis of engineered strains was done with plate reader assay. In brief, a single colony of the recombinant yeast strains was placed in 10 mL of SC medium and cultured at 30°C with shaking at 220 rpm for 24 h. 200 μL aliquot of the appropriately diluted cell suspension was taken and the cell density (OD_600_) and fluorescence intensity (excitation wavelength at 488nm, emission wavelength at 509 nm) were measured on a microplate reader. The mean fluorescence intensity was calculated by dividing the fluorescence intensity by the cell density.

For dynamic fluorescence analysis of the engineered sensor strains, a single colony of sensor strain was inoculated into 3 mL SC medium and grown at 30 °C with shaking at 200 rpm to an OD_600_ between 0.6 and 0.8. 100 μL aliquot of the cell suspension was added to a 96-well black clear-bottom plates (Corning Costar, Corning, NY, USA), followed by the addition of 1 μL of dioxin-like compounds (β-NF dissolved in DMSO, TCDD in DMSO, BaP in methanol, PCB-126 in isooctane, PCB-169 in isooctane) in a gradient concentration range from 1×10^−9^ M to 1×10^−5^ M, repeated in triplicate. Before adding the sensor strains, the solvents of the compounds BaP, PCB-169, and PCB-126 were completely evaporated on a clean bench to minimize solvent toxicity. 1 μL DMSO, with a concentration lower than 1% (v/v) was used as negative control. Afterwards, the 96-well plate was covered with a breathable sealing film (sterile sealing films; Axygen) and placed in a paradigm detection platform (Beckman Coulter) plate reader for signal detection and data acquisition. The fluorescence signal was read every 15 minutes for 8 h, and the microplate was continuously randomly shaken at 30 °C to maintain optimal signal output.

### Data and statistical analysis

Dose-response data were obtained by measuring the fluorescence output of the biosensors to different concentrations of the target analyte in a 96-well plate. The maximum fluorescence values from the kinetic readings for each well in the 96-well plate were fitted to a Hill function model by nonlinear regression (GraphPad Prism, Version 9.4.1, GraphPad Software Inc., San Diego, CA, USA). Key parameters such as Hill constant (*Kc*), Hill coefficient (*n*), and maximum fluorescence output (*k*) were obtained from sigmoidal curves. The fold change, expressed as the ratio of the mean fluorescence signal of technical triplicate wells to that of the negative control (DMSO as stimulant), was calculated for each analyte concentration in individual assay. The mean and standard deviation were calculated from the replicate readings to determine the variability between assays.

### Calculation of limit of detection (LOD), noise factor (NF), and signal-to-noise ratio (SNR)

As previously reported (33), the detection limit is defined as the lowest concentration of analyte that produces a signal that can be reliably distinguished from the “analyte noise”, at which detection is possible. Analytical noise refers to the signal observed in the absence of analyte, and is described as the limit of blank (LOB).

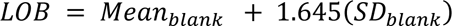

where LOB is estimated by measuring replicates of the blank sample to determine the mean and standard deviation (SD) in the absence of the analyte. Assuming that the raw analytical data from the blank samples is Gaussian distributed, the LOB covers 95% of the observed values. The LOD is determined by considering the estimated LOB value and the measurement results of replicate samples known to contain low concentrations of the analyte. It can be described as

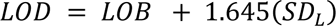

Where SD_L_ represents the standard deviation calculated from the measurement replicates of samples containing a low concentration of the analyte.

The noise factor (NF) and signal-to-noise ratio (SNR) were calculated as reported in a previous study (11). Specifically, the noise factor of the sensor is the ratio of the SNR at the input (SNR_I_, the initial sensor output) divided by the SNR at the output (SNR_O,_ the sensor output after amplification): NF = SNR_I_ /SNR_O_. The SNR was calculated by taking the difference between the averaged induced and uninduced sensor outputs, dividing it by the standard deviation of the processed sensor outputs (34).

## Results and Discussion

### Design and construction of biosensors for DLC detection

As a demonstration, we first constructed a WCB based on the aryl hydrocarbon receptor (AhR) for the detection of doxin-like chemicals (DLCs). The AhR signaling pathway depicted in **Fig. S1** shows that AhR, a ligand-dependent transcription factor, acts as a molecular switch in response to DLCs. The N-terminal nuclear localization sequence (NLS) of AHR was first exposed, causing the DLCs-AHR complex to translocate to the nucleus upon binding DLCs. Subsequently, the AHR bound to DLCs undergoes heterodimerization with its nuclear translocator ARNT to form a ternary DLCs–AHR–ARNT complex, which is further recruited to the dioxin response element (DRE, a common DNA consensus motif, 5′-TNGCGTG-3′, thereby driving the transcription of downstream genes (22). Therefore, a prototype biosensor for DLCs response was constructed in *S. cerevisiae*, which was simplified into a two-stage processor, including a sensing module that recognizes and converts external signals and a report module that generates a detectable fluorescent signal **(Fig. 1a)**.

**Figure 1.**
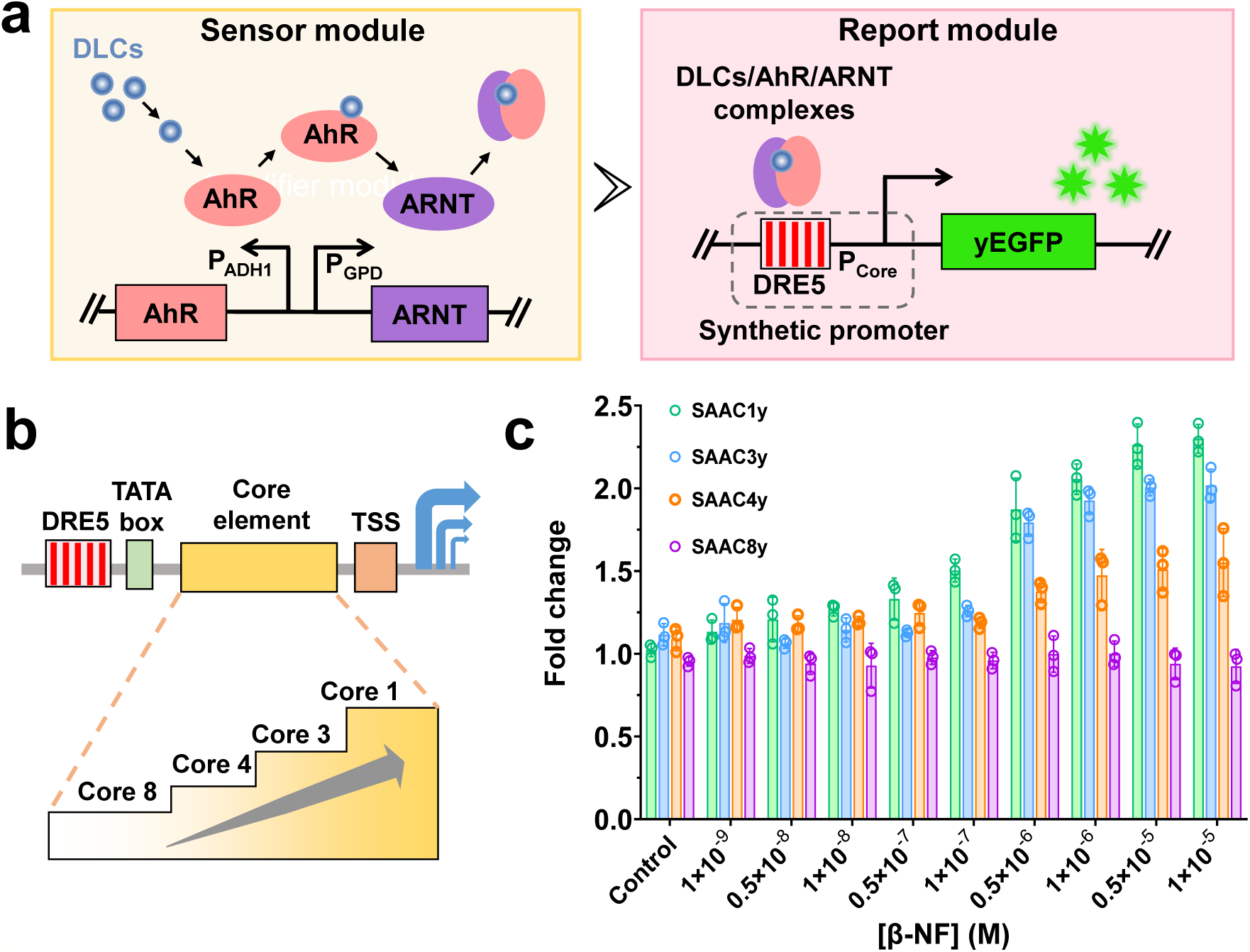
Construction and characterization of engineered yeast biosensors for DLC response. **a,** Schematic diagram of the prototype yeast sensor for DLC response, including a sensor module for recognition of inducers and a report module for generating fluorescent signals. DLCs, Doxin-like chemicals. AhR, aryl hydrocarbon receptor. ARNT, AHR nuclear translocator. DRE5, five dioxin response element fragments. P_Core_, the core element of synthetic promoters. yEGFP, yeast-enhanced green fluorescent protein. **b,** Optimization of synthetic promoters consisting of DRE5, TATA box, core element, and TSS by adjusting the strength of core elements. DRE5, TATA box, a region rich in thymidine and adenosine. TSS, a transcription start site. **c,** Responses of engineered yeast sensor to β-NF stimulation from synthetic promoters. Output signal of yEGFP was collected after 7 hours of cultivation. Error bars are mean ± SD (n = 3 biologically independent samples)

To examine the performance of the sensor in response to DLC stimulation, we introduced two yeast episomal plasmids, pESD-AhR-ARNT (CEN/ARS *ori*, AhR and ARNT are constitutive expressed under P_ADH1_ and P_GPD_ promoter, respectively) and pBEVY-DRE5-P_Core_-yEGFP (2μ *ori*, yEGFP is expressed under the synthetic promoter DRE5-P_Core_ activated by DLCs–AHR–ARNT complexes) **(Table S1)**. The synthetic promoter DRE5-P_Core_ is a homologous driving promoters of the DLCs–AHR–ARNT complexes and was specifically developed based on the minimal synthetic yeast promoters reported by Redden *et al.* (30). The effect of synthetic promoter on the sensor capacity was studied through selecting four core elements (with different strength: P_Core1_ > P_Core3_ > P_Core4_ > P_Core8_) to investigate the effect of the synthetic promoter on the sensor capacity **(Fig. 1b)**. We co-transformed these two plasmids into *S. cerevisiae* EBY100 (31) and obtained four distinct transformants, namely SAAC1y, SAAC3y, SAAC4y, and SAAC8y. The low-toxic AhR agonist β-naphthoflavone (β-NF) was provided in a gradient concentration range from 1×10^−9^ M to 1×10^−5^ M. The dynamic fluorescence properties of SAAC1y, SAAC3y, SAAC4y show that the fluorescence intensity increases with the extension of induction time. The signal appears after about 2.5 hours of cultivation and reached a peak at about 7 hours **(Fig. S2)**, except for SAAC8y. That is, synthetic promoter strengths incorporated with weak regulatory elements may result in low dynamic range output. During the assay, the fold change (maximum fluorescence intensity compared to background) not only increased linearly with the detectable inducer concentrations, but also positively correlated with the intensity of the core elements. Thus, for SAAC1y, the maximal fold change occurs at 2.3-fold after 1×10^−5^ M β-NF treatment, and the detection limit was calculated to be 4.59×10^−8^ M **(Fig. 1c, Table S4)**. These findings demonstrate for the first time that yeast sensors based on the yEGFP reporter gene are able to respond to DLCs.

### Developing a transcriptional activation component to improve the performance of the yeast sensor

To enhance the performance of the prototype yeast sensor, we attempted to develop an amplification module that processes the signal obtained from sensor module and then converts the signal input to the report module **(Fig. 2c)**. To this end, we used a series of pathway-specific regulatory proteins involved in the yeast *GAL* genetic network to verify the feasibility of our design **(Fig. S3)** (25). In fact, Blank *et al*. have reported that the transcription factor Gal4 and Gal3 regulator and its three mutants (F237Y, D368V, and S509P) are able to initiate transcriptional activation of the Gal4 protein in the absence of galactose (35), so we selected them as potential candidates for signal amplifiers.

**Figure 2.**
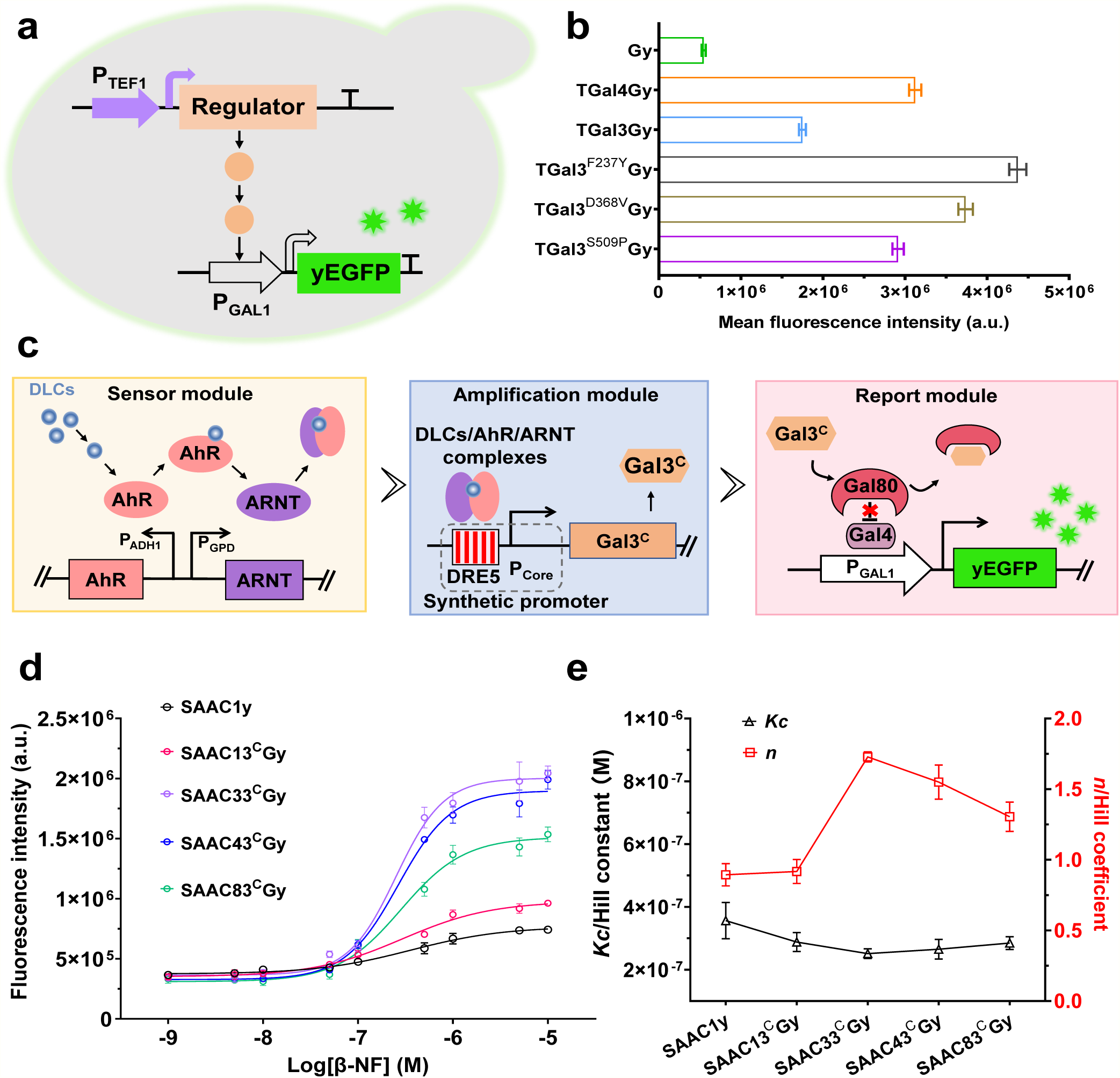
Engineering an amplifier-based biosensor with adjustable performance characteristics. **a,** A regulatory pathway was developed for screening regulatory proteins capable of activating the P_GAL1_ promoter in glucose-containing cultivation. Two expression cassettes were designed to carry out the screening task, one for constitutively expressed regulator proteins under the P_TEF1_ promoter and the other for expressing the yEGFP reporter under the P_GAL1_ promoter activated by the regulator. **b,** The mean fluorescence intensity (fluorescence intensity/OD_600_) of cells integrated with regulatory pathway and equipped with the transcription factor Gal4, the Gal3 regulator and its three mutants (F237Y, D368V, and S509P), respectively. Gy is a transformant containing only the yEGFP reporter expression cassette and was used as a control. Values are mean ± s.d. (n = 3 biologically independent experiments). **c,** Architecture of amplifier-based biosensor for DLCs response, including sensor module (recognition and processing of DLCs), amplification module (conversion and amplification of signals), report module (emission of fluorescent signals). **d**, Effect of the synthetic promoters on the responsiveness of amplifier-based biosensor toward β-NF. The fluorescence intensity corresponding to β-NF concentration collected after 5.5 hours of cultivation, was fitted as a nonlinear sigmoidal curve. Data are presented as mean ± SD for three independent experiments. **e**, Plots illustrating the Hill constant (*Kc*) and Hill coefficient (*n*) of the sensor response to different analyte concentrations. Values are means with 95% confidence intervals.

To examine the capability of regulatory proteins on GAL promoter activation under glucose cultivation, we developed an engineered system comprising two yeast episomal plasmids: pESD-P_TEF1_-Regulator (Gal4, Gal3, Gal3^F237Y^, Gal3^D368V^, and Gal3^S509P^) and pBEVY-P_GAL1_-yEGFP, where the expression of yEGFP is initiated by P_GAL1_ promoter, and the activity of which is exclusively controlled by regulatory proteins **(Fig. 2a)**. We then co-transformed the two plasmids into *S. cerevisiae* EBY100 to obtain five recombinant strains TGal4Gy, TGal3Gy, TGal3^F237Y^Gy, TGal3^D368V^Gy, TGal3^S509P^Gy, whereas the recombinant strains Gy transformed with pBEVY-P_GAL1_-yEGFP alone was used as a control strain. After incubation in glucose-containing medium for 24 h, 200 μL samples were taken to determine the mean fluorescence intensity. As shown in **Fig. 2b**, TGal3^F237Y^Gy exhibits the most significant mean fluorescence intensity among the tested strains. These data suggest that Gal3^F237Y^ can be envisioned as a tunable amplifier candidate, which helps to improve the overall performance of the sensor we designed. For ease of description, we renamed Gal3^F237Y^ mutant as Gal3^C^, following the terminology used by Blank *et al.* (35)

As a proof of concept, we developed an amplifaction module between sensor module and report module, in which the Gal3^C^ amplification is mediated by the synthetic promoter DRE5-P_Core_ and controlled by the DLCs–AHR–ARNT complexes **(Fig. 2c)**. Given that the performance of the yeast sensor is closely related to the relative concentration of Gal3^C^, and that varying the strength of the DRE5-P_Core_ promoter is a straightforward and predictable solution, we also characterized the four synthetic promoters on the sensor amplification effect. For ease of operation, we integrated the Gal3^C^ expression cassette into the reporter module to generate pBEVY-Gal3^C^-P_Core_-DRE5-P_GAL1_-yEGFP. After co-transformation with pESD-AhR-ARNT into EBY100 competent cell, the transformants were named SAAC13^C^Gy, SAAC33^C^Gy, SAAC43^C^Gy, and SAAC83^C^Gy. Under β-NF stimulation, the dynamic fluorescence properties showed that the fluorescence intensity of SAAC33CGy, SAAC43CGy, and SAAC83CGy increased with the extension of induction time, the signal appeared after approximately 1.5 hours of cultivation and reached a peak value around 5.5 hours **(Fig. S4b, c, d)**, and the response time was significantly shortened compared to SAAC1y. Although the synthetic promoter DRE5-P_Core1_ containing a strong core element in SAAC13^C^Gy, the response ability is weak **(Fig. S4a)**. We speculate that high intracellular concentrations of Gal3^C^ may exert an inhibitory effect on the transcriptional activation of the P_GAL1_ promoter.

We further implemented a Hill function-based biochemical method to fit its dose-response curve **(Fig. 2d)**, and found that the SAAC33^C^Gy exhibited superior response behavior to comparable sensors with integrated amplifaction modules. As shown in **Fig. 2e** and **Table S4**, compared to SAAC1y, SAAC33^C^Gy shows a 1.9-fold increase in sensitivity (*n*), a 1.4-fold increase in the LOD, and a 1.4-fold increase in half-maximum effective concentration (EC_50_, *Kc*). Most importantly, the maximum fluorescence output (*k*) of SAAC33^C^Gy is approximately 2.8 times that of SAAC1y. Moreover, the low noise factor (generally less than 2.0) demonstrated that no substantial noise was introduced during the signal amplification stage (11). These data clearly show that the Gal3^C^-base amplification module can be rewired into the biosensor to adjust the response characteristic curve.

### CRISPR-mediated transcriptional recruitment module for enhancing the function of Gal3^C^-base amplifier

To meet the need of future point-of-care detection, improving the sensitivity and output magnitude of biosensor remains pivotal for optimizing biosensors (19,36). Due to the integration of various regulatory frameworks, a multi-level platform will be formed in which each modality can find its niche and cooperate to execute complex functions, and is widely used in synthetic genetic circuits for multi-level signal processing (37). We next attempted to use the CRISPR-dCas9 technology capable of genome-wide regulation (38,39) as another regulatory system in this multi-level architecture, in addition to Gal3^C^-based signal amplification, to further optimize the performance of our engineered yeast sensor. Considering that dCas9 fused to a single effector domain generally has a weak transcriptional regulatory effect, and the transcriptional output depends largely on the number of transcriptional effectors recruited to the promoter (20,40), we combined the protein coupling technology SpyTag/SpyCatcher (abbreviated as SpyT/SpyC) (27) with CRISPR–dCas9 technology to enhance the transcriptional activation effect. As shown in **Fig. 3a**, we designed an optimized signal amplifier including a transcriptional activation component promoted by the Gal3^C^ regulator and a transcription recruitment component regulated by the dCas9-(SpyC)_n_ protein complex, collectively termed as a synergistic amplifier. In theory, our synergistic amplifier has the versatility of dCas9-SpyC and its ability to regulate DNA targeting and/or effector recruitment, and thus can significantly improve the signal output of biosensors.

**Figure 3.**
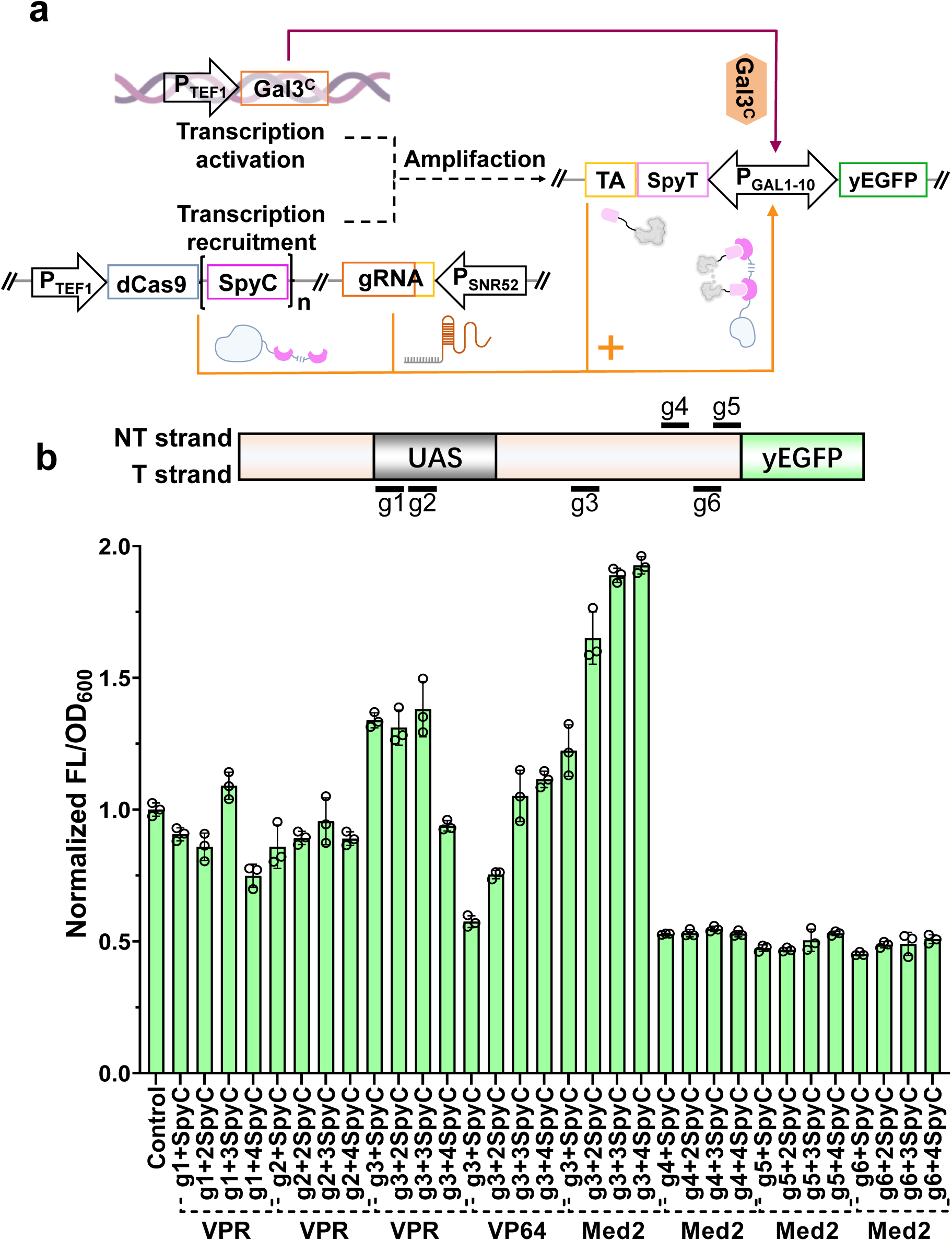
Proof-of-concept for enhanced reporter signal output involving coupling of transcriptional activation components and transcriptional recruitment components. **a**. Schematic diagram of a genetic circuit to improve yEGFP output by integrating the following components: The transcriptional activation component of the Gal3^C^ regulator is mediated by the constitutive promoter P_TEF1_; The transcription recruitment component containing the dCas9-(SpyC)_n_ fusion and guide RNA (gRNA) are constitutively expressed under the P_TEF1_ and P_SNR52_ promoters, respectively; The signal-producing component including yEGFP and TA-SpyT fusion are simultaneously controlled by the P_GAL1-10_ hybrid bidirectional promoter. TA, transcription activator. **b.** Activity analysis of genetic circuits through optimizing the regulatory capabilities of transcriptional activators (VPR, VP64, Med2) and gRNA target regions. Upper panel shows the region of DNA sequences targeted by gRNA. T strand, template strand. NT strand, non-template strand. Lower panel shows that yEGFP fluorescence is quantified by a microplate reader and displayed as mean fluorescence intensity, with data normalized to the activity of the control strain, where the uncoupled transcriptional recruitment component was used as the control strain. Data are displayed as mean ± SD for three independent experiments.

Thus, a genetic circuit was constructed containing a transcriptional activation component (yeast integrative plasmid YIPlac204-P_TEF1_-Gal3^C^, where Gal3^C^ regulator activation was mediated by the constitutive promoter P_TEF1_), a transcriptional recruitment component (yeast episomal plasmid pRS425-dCas9-(SpyC)_n_-gRNA, where the transcription recruitment element dCas9-(SpyC)_n_ and guide RNA (gRNA) were constitutively expressed under the P_TEF1_ and P_SNR52_ promoters, respectively) and a signal output component (yeast episomal plasmid pBEVY-TA-SpyT-P_GAL1-10_-yEGFP, where the P_GAL1-10_ hybrid bidirectional promoter was selected to simultaneously express yEGFP and TA-SpyT fusion. TA: abbreviation for transcriptional activator). In principle, the expression of TA-SpyT fusion protein and yEGFP are both regulated simultaneously by the Gal3^C^-activated P_GAL1-10_ hybrid bidirectional promoter. Once TA-SpyT is produced, it will be captured by dCas9-(SpyC)_n_ through SpyC-SpyT conjugation. Subsequently, under the combined action of SpyC gene dosage and gRNA guidance, a higher local concentration of TA-SpyT is targeted in the pre-defined DNA sequence of P_GAL1-10_ in a positive feedback manner, ultimately leading to further enhancement of yEGFP output **(Fig. 3a)**. To confirm that coupling transcriptional activation and transcriptional recruitment components cumulatively enhances the output of signals, we optimized the activity of the genetic circuit by adjusting the regulatory capacity of the transcription activator (VPR, VP64-p65-Rta tripartite activator (41); VP64: tetrameric repeats of the herpes simplex virus protein VP16 (26); Med2: a subunit of the RNA polymerase II mediator complex (20)) and the target regions of gRNA. As depicted in **Fig. 3b**, when g3 was used as the gRNA and VPR was used as the transcriptional activator, we observed an approximately 1.4-fold increase in the mean fluorescence intensity output from the genetic circuit. In addition, under the guidance of g3, the activation efficiency is the highest, which increased by nearly 2-fold, and the mean fluorescence intensity of yEGFP increased almost linearly with the number of SpyC copies. Further evaluation of the DNA sequences targeted by gRNA also found that among g3-g6, g3 yielded a remarkable reporter signal output **(Fig. 3b)**. These data strongly suggest that high levels of reporter signal output can be achieved through a custom engineering process involving the coupling of the transcriptional activation component and the transcription recruitment component, which can serve as a prerequisite for guiding rational amplifier efforts to provide better sensitivity and output amplitude.

### The synergistic effect of transcriptional activation and transcriptional recruitment confers benefits on sensor behaviors

Based on the characteristics of the synergistic effect of transcriptional activation and transcriptional recruitment, a synergistic amplification module between the sensor module and the reporter module was developed, as shown in **Fig. S5**. However, we found that yeast sensor with integrated synergistic amplifiers show little response to β-NF stimulation (data not shown). We speculate that this phenomenon, in the sensing pathway, is caused by bidirectional regulation of P_GAL1-10_, whose transcriptional activation ability is reduced compared to P_GAL1_ promoted by Gal3^C^ regulator. We then refined the synergistic amplification module based on the polycistronic expression regulated by the intergenic sequence of *IGG6* (32). As shown in **Fig. 4a**, a bicistronic cassette containing the *Gal3^C^* and *Med2-SpyT* genes downstream of the synthetic promoter was developed, which possesses the ability to simultaneously trigger the expression of genes involved in transcriptional activation and transcriptional recruitment, and allows the initiation of yEGFP expression via P_GAL1_ but not P_GAL1-10_.

**Figure 4.**
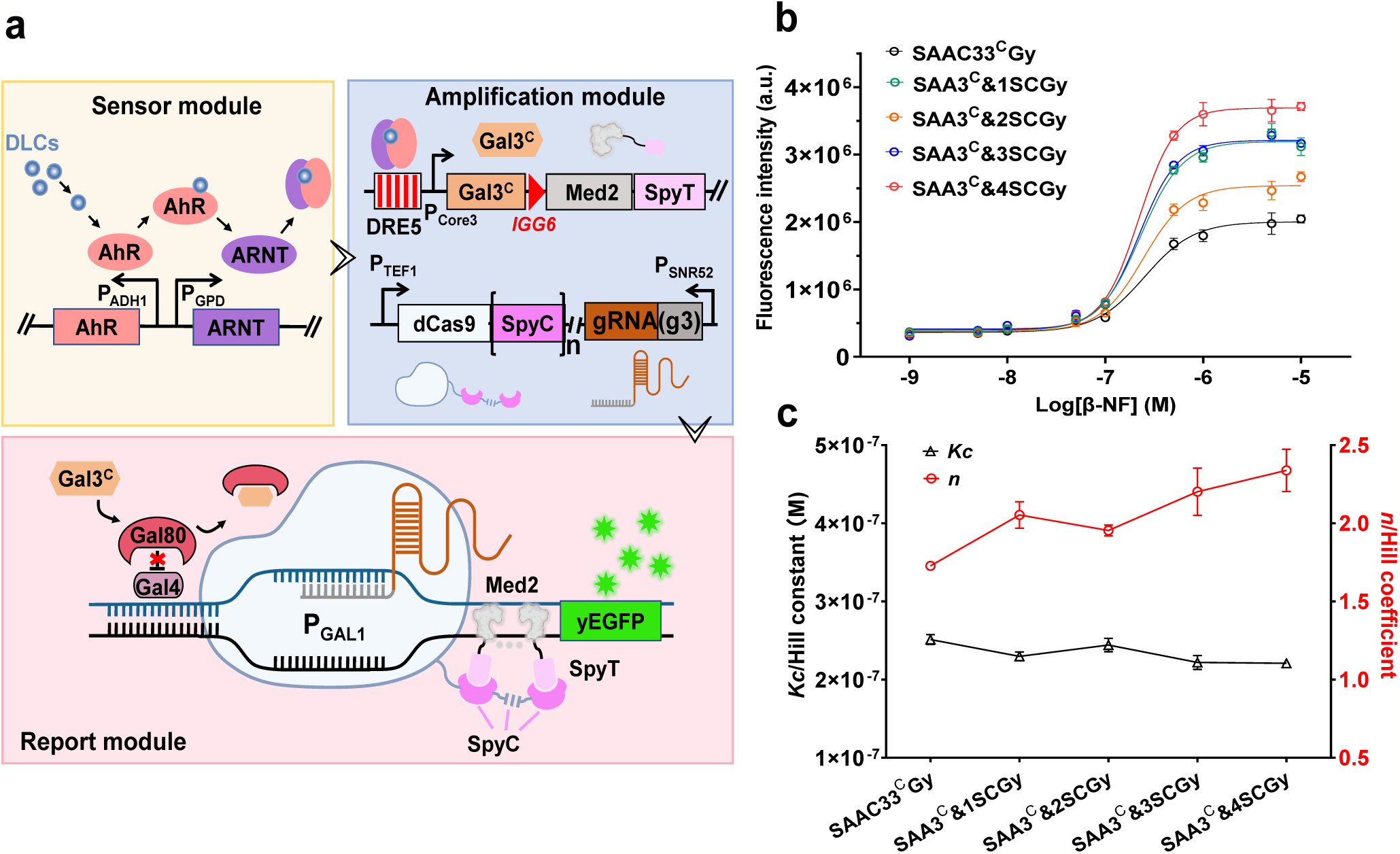
Developing a synergistic amplifier to augment the inherent sensing capabilities of yeast biosensors. **a,** Schematic diagram illustrating the synergistic amplifier-based biosensor for DLCs response, comprising a transcriptional activation component carried out by Gal3^C^ regulator and a transcriptional recruitment component regulated by dCas9-(SpyC)_n_ protein complex, together termed as a synergistic amplifier. *IGG6*, an intergenic sequence that functions to trigger the genes of Ga*l3^C^* and *Med2-SpyT* expression simultaneously. **b**, Dose-response curves of the yeast sensors integrated with multiple tandem repeat SpyC units in the synergistic amplifier following β-NF stimulation, SAA3^C^&1SCGy, SAA3^C^&2SCGy, SAA3^C^&3SCGy, SAA3^C^&4SCGy indicate yeast biosensors introduced with the synergistic amplifier, among which transcriptional recruitment component of corresponding sensor is equipped with 1, 2, 3, and 4 copies of SpyC, respectively. SAAC33^C^Gy served as a control. Data represent as mean ± SD for three independent experiments. **c**, Plots showing the Hill constant (*Kc*) and the Hill coefficient (*n*) of the sensor’s dose-response. Values are means with 95% confidence intervals.

To elucidate the extent of coupling of transcriptional activation and transcriptional recruitment on signal amplification, two plasmids, pESD-AhR-ARNT-dCas9-(SpyC)_n_-g3 and pBEVY-SpyT-Med2-*IGG6*-Gal3^C^-P_Core3_-DRE5-P_GAL1_-yEGFP, were constructed. After co-transformation into EBY100 competent cell, the transformants were named SAA3^C^&1SCGy, SAA3^C^&2SCGy, SAA3^C^&3SCGy, SAA3^C^&4SCGy, respectively. Upon β-NF stimulation, the dynamic fluorescence properties showed that the fluorescence intensity of the four sensors increased over induction time. The signal appeared after approximately 1.5 hours of cultivation and peaked at approximately 6 hours **(Fig. S6)**, similar to the signal observed for SAAC33^C^Gy. However, the combination of transcriptional activation and transcriptional recruitment results in continued improvements in sensor output and sensitivity. We can conclude that SC33^C^Md4SGy achieved the best performance with a 1.4-fold and 2.6-fold higher sensitivity than SAAC33^C^Gy and SAAC1y, respectively, based on dose-response curve. Moreover, compared to SAAC33^C^Gy and SAAC1y, the *Kc* of the synergistic amplifier-based biosensor was improved by 1.2-fold and 1.6-fold, respectively, and *k* was significantly improved by 1.8-fold and 5.1-fold respectively **(Fig. 4c, Table S4)**. Low noise coefficients (generally less than 2.0) were also observed in engineered sensors during signal amplification using a synergistic amplifier. We did notice that for SAA3^C^&2SCGy, SAA3^C^&3SCGy, SAA3^C^&4SCGy (except for SAA3^C^&1SCGy), the output dynamic range increased almost linearly with the number of SpyC copies. Therefore, we speculate that this phenomenon may be attributed to a dynamic imbalance between transcriptional activation and transcriptional recruitment. In other words, higher concentrations of transcriptional activator may hinder Gal3^C^ regulation, but this effect can be alleviated by enhancing transcriptional activator recruitment.

### Evaluation of the response effects of the synergistic amplifier to various dioxin analogues

Encouraged by the synergistic effects of transcriptional activation and transcriptional recruitment, we believe that our synergistic amplifier will tremendously enhance the potential for the practical application of yeast sensors in response to other dioxin analogues, which can also activate AhR and its downstream gene expression, but with different potencies (23). Therefore, we selected four dioxin analogues, 2,3,7,8-tetrachlorodibenzo-p-dioxin (TCDD), benzo[a]pyrene (BaP), 3,3’,4,4’,5-pentachlorobiphenyl (PCB-126), and 3,3’,4,4’,5,5’-hexachlorobiphenyl (PCB-169), as persistent toxic chemicals associated with health impacts, to determine the responsiveness of our engineered yeast sensors. We observed that our sensors exhibited enhanced response after stimulation with four dioxin analogues, showing the sensing ability in the order of SAA3^C^&4SCGy > SAAC33^C^Gy > SAAC1y. Compare to SAAC1y, the maximum fluorescence output of SAA3^C^&4SCGy increased by 25%, 40%, 48%, and 16% when treated with TCDD, Bap, PCB-126, and PCB-169, respectively, but only increased by 15%, 12%, 29%, and 7% for SAAC33^C^Gy (**Fig. 5**). Moreover, the synergistic amplifier has the ability to modulate the operating range, resulting in a sustained and significant signal output even under low-dose AhR agonist stimulation, especially at 1×10^−8^ M, thereby enhancing the detection limit (**Fig. 5**). We did notice that a subtle signal amplification was observed under the condition of PCB-169 stimulation (**Fig. 5d**). This may be attributed to its reduced accessibility to yeast sensors under aqueous culture conditions. Indeed, it has been found that increasing the number of chlorine atoms leads to an increase in hydrophobicity, a phenomenon that has been documented previously (23,28). Collectively, these findings demonstrate that the inherent ability of yeast biosensors to respond to environmental contaminants can be enhanced through the synergistic amplifiers.

**Figure 5.**
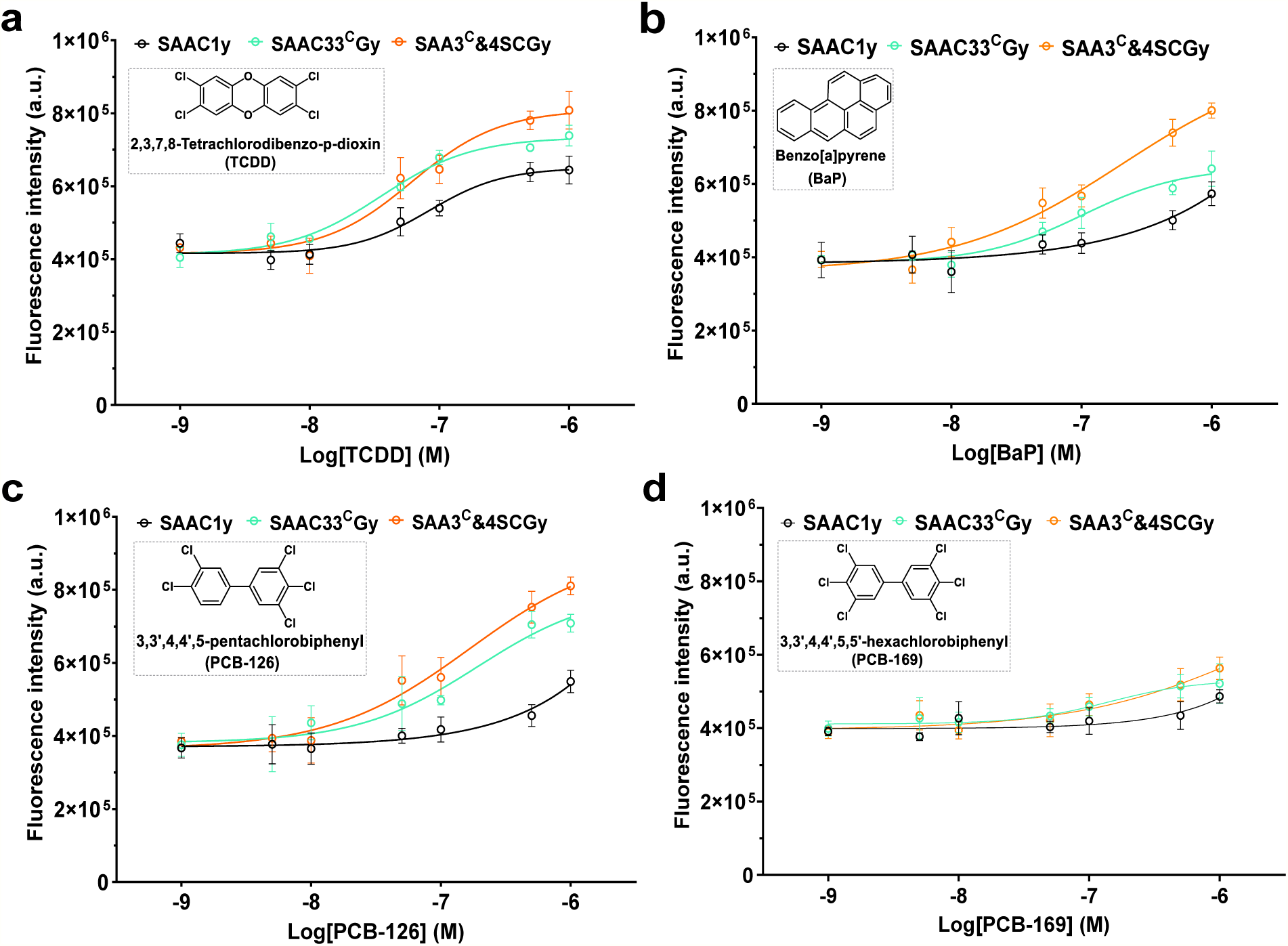
Detection of dioxin homologues with engineered sensor. Dose-response curves of the engineered yeast sensors exposed to 2,3,7,8-tetrachlorodibenzo-p-dioxin (TCDD, **a**), benzo[a]pyrene (BaP, **b**), 3,3’,4,4’,5-pentachlorobiphenyl (PCB-126, **c**), and 3,3’,4,4’,5,5’-hexachlorobiphenyl (PCB-169, **d**) under similar experimental conditions. The sensing cell were exposed to dioxin analogues at a final concentration ranging from 1×10^−9^ M to 1×10^−6^ M and cultured for 8 hours, and the fluorescence signal at the peak intensity was fitted into a nonlinear sigmoidal curve. Values represent the means ± standard deviation of triplicate determinations.

Due to the modularity and orthogonality of Gal3^C^ regulator and dCas9-(SpyC)_n_ fusion, they can be introduced individually or synergistically into various gene regulatory networks to systematically amplify and modulate signal transduction processes to achieve a wide range of biotechnological applications (42,43). For instance, the predictable adjustment in input signals facilitate the interfacing and matching of genetic modules to a broad range of dynamic response characteristics, enabling precise regulation of gene expression to fine-tune metabolite levels in synthetic cellular metabolic pathways. Although the synergistic amplifier opens up new opportunities for signal amplification, there are still some obstacles that need to be addressed. In this study, the amplifier was used only to detect DLCs. More sensors or synthetic genetic circuits are needed to adapt to diverse scenarios, making it more adaptable and versatile. Moreover, further work should be devoted to obtaining shorter response times to compete with analytical techniques such as high-performance liquid chromatography (HPLC).

## Conclusions

In summary, a novel modular signal amplification method termed synergistic amplifier was developed to substantially enhance the performance of yeast biosensors in response to dioxin-like compounds (DLCs) in terms of maximum output, detection limit, and sensitivity. We demonstrate that the synergistic amplifier is consisted of two components, a transcriptional activation component mediated by the Gal3^C^ regulator and a transcriptional recruitment component that exploits dCas9-(SpyC)_n_ function, which work together synergistically to re-program the sensor behavior. The characteristics of the synergistic amplifier can be tuned by optimizing the strength of synthetic promoter, regulatory proteins for transcriptional regulation, the number of SpyC repeats, transcriptional activators, and the gRNA target region. By using various dioxin homologues to evaluate the potential for predictive modulation, we observed progress in both the dynamic output and detection limit of the engineered sensors. This work shows that the synergistic amplifier can be integrated into biosensor architectures to reprogram their behaviors, providing a new paradigm for expanding signal processing pathways within synthetic gene networks.

## Data availability

The data underlying this article are available in the article and in its online supplementary material

## Supplementary Data

Supplementary Data are available at NAR Online

## Author contributions

CD performed the main experiments, data curation, and drafted the manuscript. YW performed data curation. JW, SL, YZ, and LY did formal analysis, Data curation. JG critically revised the manuscript, GZ designed, funded the study, and critically revised the manuscript, and all authors contributed to the writing, reading, and approval of the final manuscript.

## Declarations

The authors declare that they have no conflict of interest or personal relationships in this paper.

