## Supplementary information for "Synergistic amplifier reprograms biosensor behavior to facilitate detection of dioxin-like compounds"

**Table S1 Plasmids used in this study**

| Plasmid | Description | Reference |
|---|---|---|
| YIPlac204 | Yeast integration plasmid, LEU2 selection marker, parental plasmid | Lab store |
| pESD | Yeast episomal plasmid, CEN/ARS ori, TRP1 selection marker, parental plasmid | Lab store |
| pBEVY | Yeast episomal plasmid, 2μ ori, LEU2 or URA3 selection marker, parental plasmid | Lab store |
| pRS425 | Yeast episomal plasmid, 2μ ori, TRP1 selection marker, parental plasmid | Lab store |
| pESD-AhR-ARNT | pESD carrying *AhR* and *ARNT* gene expression cassettes, TRP1 selection marker | This study |
| pBEVY-DRE5-$P_{Core1}$-yEGFP | pBEVY carrying DRE5-$P_{Core1}$-yEGFP expression cassette, LEU2 selection marker | This study |
| pBEVY-DRE5-$P_{Core3}$-yEGFP | pBEVY carrying DRE5-$P_{Core3}$-yEGFP expression cassette, LEU2 selection marker | This study |
| pBEVY-DRE5-$P_{Core4}$-yEGFP | pBEVY carrying DRE5-$P_{Core4}$-yEGFP expression cassette, LEU2 selection marker | This study |
| pBEVY-DRE5-$P_{Core8}$-yEGFP | pBEVY carrying DRE5-$P_{Core8}$-yEGFP expression cassette, LEU2 selection marker | This study |
| pESD-$P_{TEF1}$-Gal4 | pESD carrying $P_{TEF1}$-Gal4 expression cassette, TRP1 selection marker | This study |
| pESD-$P_{TEF1}$-Gal3 | pESD carrying $P_{TEF1}$-Gal3 expression cassette, TRP1 selection marker | This study |
| pESD-$P_{TEF1}$-Gal3$^{F237Y}$ | pESD carrying $P_{TEF1}$-Gal3$^{F237Y}$ expression cassette, TRP1 selection marker | This study |
| pESD-$P_{TEF1}$- Gal3$^{D368V}$ | pESD carrying $P_{TEF1}$-Gal3$^{D368V}$ expression cassette, TRP1 selection marker | This study |
| pESD-$P_{TEF1}$- Gal3$^{S509P}$ | pESD carrying $P_{TEF1}$-Gal3$^{S509P}$ expression cassette, TRP1 selection marker | This study |
| pBEVY-$P_{GAL1}$-yEGFP | pBEVY carrying $P_{GAL1}$-yEGFP expression cassette, LEU2 selection marker | This study |
| pBEVY-Gal3$^{C}$-$P_{Core1}$-DRE5-$P_{GAL1}$-yEGFP | pBEVY carrying DRE5-$P_{Core1}$-Gal3$^{C}$ and $P_{GAL1}$-yEGFP expression cassettes, LEU2 selection marker | This study |
| pBEVY-Gal3$^{C}$-$P_{Core3}$-DRE5-$P_{GAL1}$-yEGFP | pBEVY carrying DRE5-$P_{Core3}$-Gal3$^{C}$ and $P_{GAL1}$-yEGFP expression cassettes, LEU2 selection marker | This study |

| | | |
|---|---|---|
| pBEVY-Gal3$^{C}$-P$_{Core4}$-DRE5-P$_{GAL1}$-yEGFP | pBEVY carrying DRE5-P$_{Core4}$-Gal3$^{C}$ and P$_{GAL1}$-yEGFP expression cassettes, LEU2 selection marker | This study |
| pBEVY-Gal3$^{C}$-P$_{Core8}$-DRE5-P$_{GAL1}$-yEGFP | pBEVY carrying DRE5-P$_{Core8}$-Gal3$^{C}$ and P$_{GAL1}$-yEGFP expression cassettes, LEU2 selection marker | This study |
| YIPlac204-P$_{TEF1}$-Gal3$^{C}$ | YIPlac204 carrying P$_{TEF1}$-Gal3$^{C}$ expression cassette, LEU2 selection marker | This study |
| pRS425-dCas9-SpyC-gRNA (g1-g6) | pRS425 carrying P$_{TEF1}$-dCas9-SpyC and P$_{SNR52}$-gRNA (g1-g6) expression cassettes, TRP1 selection marker | This study |
| pRS425-dCas9-(SpyC)$_2$-gRNA (g1-g6) | pRS425 carrying P$_{TEF1}$-dCas9-(SpyC)$_2$ and P$_{SNR52}$-gRNA (g1-g6) expression cassettes, TRP1 selection marker | This study |
| pRS425-dCas9-(SpyC)$_3$-gRNA (g1-g6) | pRS425 carrying P$_{TEF1}$-dCas9-(SpyC)$_3$ and P$_{SNR52}$-gRNA (g1-g6) expression cassettes, TRP1 selection marker | This study |
| pRS425-dCas9-(SpyC)$_4$-gRNA (g1-g6) | pRS425 carrying P$_{TEF1}$-dCas9-(SpyC)$_4$ and P$_{SNR52}$-gRNA (g1-g6) expression cassettes, TRP1 selection marker | This study |
| pBEVY-VPR-SpyT-P$_{GAL1-10}$-yEGFP | pBEVY carrying VPR-SpyT-P$_{GAL1-10}$-yEGFP bidirectional expression cassettes, URA3 selection marker | This study |
| pBEVY-VP64-SpyT-P$_{GAL1-10}$-yEGFP | pBEVY carrying VP64-SpyT-P$_{GAL1-10}$-yEGFP bidirectional expression cassettes, URA3 selection marker | This study |
| pBEVY-Med2-SpyT-P$_{GAL1-10}$-yEGFP | pBEVY carrying Med2-SpyT-P$_{GAL1-10}$-yEGFP bidirectional expression cassettes, URA3 selection marker | This study |
| pESD-AhR-ARNT-dCas9-SpyC-g3 | pESD carrying *AhR* and *ARNT* gene expression cassettes, along with P$_{TEF1}$-dCas9-SpyC and P$_{SNR52}$-g3 expression cassettes, TRP1 selection marker | This study |
| pESD-AhR-ARNT-dCas9-(SpyC)$_2$-g3 | pESD carrying *AhR* and *ARNT* gene expression cassettes, along with P$_{TEF1}$-dCas9-(SpyC)$_2$ and P$_{SNR52}$-g3 expression cassettes, TRP1 selection marker | This study |
| pESD-AhR-ARNT-dCas9-(SpyC)$_3$-g3 | pESD carrying *AhR* and *ARNT* gene expression cassettes, along with P$_{TEF1}$-dCas9-(SpyC)$_3$ and P$_{SNR52}$-g3 expression cassettes, TRP1 selection marker | This study |
| pESD-AhR-ARNT-dCas9-(SpyC)$_4$-g3 | pESD carrying *AhR* and *ARNT* gene expression cassettes, along with P$_{TEF1}$-dCas9-(SpyC)$_4$ and P$_{SNR52}$-g3 expression cassettes, TRP1 selection marker | This study |

| | | |
|---|---|---|
| pBEVY-SpyT-Med2-*IGG6*-Gal3$^{C}$-$P_{Core3}$-DRE5-$P_{GAL1}$-yEGFP | pBEVY carrying DRE5-$P_{Core3}$-Gal3$^{C}$-*IGG6*- Med2-SpyT bicistronic cassettes, along with $P_{GAL1}$-yEGFP expression cassette, LEU2 selection marker | This study |

**Table S2 The gRNA spacer sequences used in this study**

| gRNA | Sequence | Target | Strand | Position | PAM |
|---|---|---|---|---|---|
| g1 | GAGGAGAGTCTTCCTTCGGA | UAS of $P_{GAL1\text{-}10}$ | T | -407 to -426 | GGG |
| g2 | GTGAAGACGAGGACGCACGG | UAS of $P_{GAL1\text{-}10}$ | T | -388 to -407 | AGG |
| g3 | GATTAATTACCCCAGAAATA | $P_{GAL1}$ of $P_{GAL1\text{-}10}$ | T | -185 to -204 | AGG |
| g4 | ACTAATACTTTCAACATTTT | $P_{GAL1}$ of $P_{GAL1\text{-}10}$ | NT | -101 to -120 | CGG |
| g5 | ACGTCAAGGAGAAAAAACCC | $P_{GAL1}$ of $P_{GAL1\text{-}10}$ | NT | -8 to -27 | CGG |
| g6 | TCCTTGACGTTAAAGTATAG | $P_{GAL1}$ of $P_{GAL1\text{-}10}$ | T | -18 to -37 | AGG |

Note: T: template strand; NT: non-template strand. Position indicates the region upstream of the yEGFP start codon.

**Table S3 Engineered strains/sensors summarized in this study**

| Strains/Sensors | Genotype | Reference |
|---|---|---|
| **Engineered strains** | - | - |
| *S. cerevisae* W303-1a | *MATa; leu2-3,112; trp1-1; can1-100; ura3-1; ade2-1; his3-11,15* | Lab store |
| *S. cerevisae* EBY100 | *MATα; ura3-52; trp1; leu2Δ1; his3Δ200; pep4::HIS3; prb1Δ1.6R; can1; GAL (pIU211: URA3)* | Lab store |
| TGal4Gy | EBY100 carrying pESD-$P_{TEF1}$-Gal4 and pBEVY-$P_{GAL1}$-yEGFP | This study |
| TGal3Gy | EBY100 carrying pESD-$P_{TEF1}$-Gal3 and pBEVY-$P_{GAL1}$-yEGFP | This study |
| TGal3$^{F237Y}$Gy | EBY100 carrying pESD-$P_{TEF1}$-Gal3$^{F237Y}$ and pBEVY-$P_{GAL1}$-yEGFP | This study |
| TGal3$^{D368V}$Gy | EBY100 carrying pESD-$P_{TEF1}$-Gal3$^{D368V}$ and pBEVY-$P_{GAL1}$-yEGFP | This study |
| TGal3$^{S509P}$Gy | EBY100 carrying pESD-$P_{TEF1}$-Gal3$^{S509P}$ and pBEVY-$P_{GAL1}$-yEGFP | This study |
| Gy | EBY100 carrying pBEVY-$P_{GAL1}$-yEGFP | This study |
| VPR(g1+SpyC) | W303-1a; *LEU2::YIPlac204-$P_{TEF1}$-Gal3$^{C}$*; carrying pRS425-dCas9-SpyC-gRNA (g1) and pBEVY-VPR-SpyT-$P_{GAL1-10}$-yEGFP | This study |
| VPR(g1+2SpyC) | W303-1a; *LEU2::YIPlac204-$P_{TEF1}$-Gal3$^{C}$*; carrying pRS425-dCas9-(SpyC)$_2$-gRNA (g1) and pBEVY-VPR-SpyT-$P_{GAL1-10}$-yEGFP | This study |
| VPR(g1+3SpyC) | W303-1a; *LEU2::YIPlac204-$P_{TEF1}$-Gal3$^{C}$*; carrying pRS425-dCas9-(SpyC)$_3$-gRNA (g1) and pBEVY-VPR-SpyT-$P_{GAL1-10}$-yEGFP | This study |
| VPR(g1+4SpyC) | W303-1a; *LEU2::YIPlac204-$P_{TEF1}$-Gal3$^{C}$*; carrying pRS425-dCas9-(SpyC)$_4$-gRNA (g1) and pBEVY-VPR-SpyT-$P_{GAL1-10}$-yEGFP | This study |
| VPR(g2+SpyC) | W303-1a; *LEU2::YIPlac204-$P_{TEF1}$-Gal3$^{C}$*; carrying pRS425-dCas9-SpyC-gRNA (g2) and pBEVY-VPR-SpyT-$P_{GAL1-10}$-yEGFP | This study |
| VPR(g2+2SpyC) | W303-1a; *LEU2::YIPlac204-$P_{TEF1}$-Gal3$^{C}$*; carrying pRS425-dCas9-(SpyC)$_2$-gRNA (g2) and pBEVY-VPR-SpyT-$P_{GAL1-10}$-yEGFP | This study |
| VPR(g2+3SpyC) | W303-1a; *LEU2::YIPlac204-$P_{TEF1}$-Gal3$^{C}$*; carrying pRS425-dCas9-(SpyC)$_3$-gRNA (g2) and pBEVY-VPR-SpyT-$P_{GAL1-10}$-yEGFP | This study |
| VPR(g2+4SpyC) | W303-1a; *LEU2::YIPlac204-$P_{TEF1}$-Gal3$^{C}$*; carrying pRS425-dCas9-(SpyC)$_4$-gRNA (g2) and pBEVY-VPR-SpyT-$P_{GAL1-10}$-yEGFP | This study |
| VPR(g3+SpyC) | W303-1a; *LEU2::YIPlac204-$P_{TEF1}$-Gal3$^{C}$*; carrying pRS425-dCas9-SpyC-gRNA (g3) and pBEVY-VPR-SpyT-$P_{GAL1-10}$-yEGFP | This study |
| VPR(g3+2SpyC) | W303-1a; *LEU2::YIPlac204-$P_{TEF1}$-Gal3$^{C}$*; carrying pRS425-dCas9-(SpyC)$_2$-gRNA (g3) and pBEVY-VPR-SpyT-$P_{GAL1-10}$-yEGFP | This study |
| VPR(g3+3SpyC) | W303-1a; *LEU2::YIPlac204-$P_{TEF1}$-Gal3$^{C}$*; carrying pRS425-dCas9-(SpyC)$_3$-gRNA (g3) and pBEVY-VPR-SpyT-$P_{GAL1-10}$-yEGFP | This study |
| VPR(g3+4SpyC) | W303-1a; *LEU2::YIPlac204-$P_{TEF1}$-Gal3$^{C}$*; carrying pRS425-dCas9-(SpyC)$_4$-gRNA (g3) and pBEVY-VPR-SpyT-$P_{GAL1-10}$-yEGFP | This study |

| | | |
|---|---|---|
| VP64(g3+SpyC) | W303-1a; *LEU2::YIPlac204-$P_{TEF1}$-Gal3$^C$*; carrying pRS425-dCas9-SpyC-gRNA (g3) and pBEVY-VP64-SpyT-$P_{GAL1-10}$-yEGFP | This study |
| VP64(g3+2SpyC) | W303-1a; *LEU2::YIPlac204-$P_{TEF1}$-Gal3$^C$*; carrying pRS425-dCas9-$(SpyC)_2$-gRNA (g3) and pBEVY-VP64-SpyT-$P_{GAL1-10}$-yEGFP | This study |
| VP64(g3+3SpyC) | W303-1a; *LEU2::YIPlac204-$P_{TEF1}$-Gal3$^C$*; carrying pRS425-dCas9-$(SpyC)_3$-gRNA (g3) and pBEVY-VP64-SpyT-$P_{GAL1-10}$-yEGFP | This study |
| VP64(g3+4SpyC) | W303-1a; *LEU2::YIPlac204-$P_{TEF1}$-Gal3$^C$*; carrying pRS425-dCas9-$(SpyC)_4$-gRNA (g3) and pBEVY-VP64-SpyT-$P_{GAL1-10}$-yEGFP | This study |
| Med2(g3+SpyC) | W303-1a; *LEU2::YIPlac204-$P_{TEF1}$-Gal3$^C$*; carrying pRS425-dCas9-SpyC-gRNA (g3) and pBEVY-Med2-SpyT-$P_{GAL1-10}$-yEGFP | This study |
| Med2(g3+2SpyC) | W303-1a; *LEU2::YIPlac204-$P_{TEF1}$-Gal3$^C$*; carrying pRS425-dCas9-$(SpyC)_2$-gRNA (g3) and pBEVY-Med2-SpyT-$P_{GAL1-10}$-yEGFP | This study |
| Med2(g3+3SpyC) | W303-1a; *LEU2::YIPlac204-$P_{TEF1}$-Gal3$^C$*; carrying pRS425-dCas9-$(SpyC)_3$-gRNA (g3) and pBEVY-Med2-SpyT-$P_{GAL1-10}$-yEGFP | This study |
| Med2(g3+4SpyC) | W303-1a; *LEU2::YIPlac204-$P_{TEF1}$-Gal3$^C$*; carrying pRS425-dCas9-$(SpyC)_4$-gRNA (g3) and pBEVY-Med2-SpyT-$P_{GAL1-10}$-yEGFP | This study |
| Med2(g4+SpyC) | W303-1a; *LEU2::YIPlac204-$P_{TEF1}$-Gal3$^C$*; carrying pRS425-dCas9-SpyC-gRNA (g4) and pBEVY-Med2-SpyT-$P_{GAL1-10}$-yEGFP | This study |
| Med2(g4+2SpyC) | W303-1a; *LEU2::YIPlac204-$P_{TEF1}$-Gal3$^C$*; carrying pRS425-dCas9-$(SpyC)_2$-gRNA (g4) and pBEVY-Med2-SpyT-$P_{GAL1-10}$-yEGFP | This study |
| Med2(g4+3SpyC) | W303-1a; *LEU2::YIPlac204-$P_{TEF1}$-Gal3$^C$*; carrying pRS425-dCas9-$(SpyC)_3$-gRNA (g4) and pBEVY-Med2-SpyT-$P_{GAL1-10}$-yEGFP | This study |
| Med2(g4+4SpyC) | W303-1a; *LEU2::YIPlac204-$P_{TEF1}$-Gal3$^C$*; carrying pRS425-dCas9-$(SpyC)_4$-gRNA (g4) and pBEVY-Med2-SpyT-$P_{GAL1-10}$-yEGFP | This study |
| Med2(g5+SpyC) | W303-1a; *LEU2::YIPlac204-$P_{TEF1}$-Gal3$^C$*; carrying pRS425-dCas9-SpyC-gRNA (g5) and pBEVY-Med2-SpyT-$P_{GAL1-10}$-yEGFP | This study |
| Med2(g5+2SpyC) | W303-1a; *LEU2::YIPlac204-$P_{TEF1}$-Gal3$^C$*; carrying pRS425-dCas9-$(SpyC)_2$-gRNA (g5) and pBEVY-Med2-SpyT-$P_{GAL1-10}$-yEGFP | This study |
| Med2(g5+3SpyC) | W303-1a; *LEU2::YIPlac204-$P_{TEF1}$-Gal3$^C$*; carrying pRS425-dCas9-$(SpyC)_3$-gRNA (g5) and pBEVY-Med2-SpyT-$P_{GAL1-10}$-yEGFP | This study |
| Med2(g5+4SpyC) | W303-1a; *LEU2::YIPlac204-$P_{TEF1}$-Gal3$^C$*; carrying pRS425-dCas9-$(SpyC)_4$-gRNA (g5) and pBEVY-Med2-SpyT-$P_{GAL1-10}$-yEGFP | This study |
| Med2(g6+SpyC) | W303-1a; *LEU2::YIPlac204-$P_{TEF1}$-Gal3$^C$*; carrying pRS425-dCas9-SpyC-gRNA (g6) and pBEVY-Med2-SpyT-$P_{GAL1-10}$-yEGFP | This study |
| Med2(g6+2SpyC) | W303-1a; *LEU2::YIPlac204-$P_{TEF1}$-Gal3$^C$*; carrying pRS425-dCas9-$(SpyC)_2$-gRNA (g6) and pBEVY-Med2-SpyT-$P_{GAL1-10}$-yEGFP | This study |

| | | |
|---|---|---|
| Med2(g6+3SpyC) | W303-1a; *LEU2::YIPlac204-$P_{TEF1}$-Gal3$^{C}$*; carrying pRS425-dCas9-$(SpyC)_3$-gRNA (g6) and pBEVY-Med2-SpyT-$P_{GAL1-10}$-yEGFP | This study |
| Med2(g6+4pyC) | W303-1a; *LEU2::YIPlac204-$P_{TEF1}$-Gal3$^{C}$*; carrying pRS425-dCas9-$(SpyC)_4$-gRNA (g6) and pBEVY-Med2-SpyT-$P_{GAL1-10}$-yEGFP | This study |
| **Sensors** | - | - |
| SAAC1y | EBY100 carrying pESD-AhR-ARNT and pBEVY-DRE5-$P_{Core1}$-yEGFP | This study |
| SAAC3y | EBY100 carrying pESD-AhR-ARNT and pBEVY-DRE5-$P_{Core3}$-yEGFP | This study |
| SAAC4y | EBY100 carrying pESD-AhR-ARNT and pBEVY-DRE5-$P_{Core4}$-yEGFP | This study |
| SAAC8y | EBY100 carrying pESD-AhR-ARNT and pBEVY-DRE5-$P_{Core8}$-yEGFP | This study |
| SAAC13$^{C}$Gy | EBY100 carrying pESD-AhR-ARNT and pBEVY-Gal3$^{C}$-$P_{Core1}$-DRE5-$P_{GAL1}$-yEGFP | This study |
| SAAC33$^{C}$Gy | EBY100 carrying pESD-AhR-ARNT and pBEVY-Gal3$^{C}$-$P_{Core3}$-DRE5-$P_{GAL1}$-yEGFP | This study |
| SAAC43$^{C}$Gy | EBY100 carrying pESD-AhR-ARNT and pBEVY-Gal3$^{C}$-$P_{Core4}$-DRE5-$P_{GAL1}$-yEGFP | This study |
| SAAC83$^{C}$Gy | EBY100 carrying pESD-AhR-ARNT and pBEVY-Gal3$^{C}$-$P_{Core8}$-DRE5-$P_{GAL1}$-yEGFP | This study |
| SAA3$^{C}$&1SCGy | EBY100 carrying pESD-AhR-ARNT-dCas9-SpyC-g3 and pBEVY-SpyT-Med2-*IGG6*-Gal3$^{C}$-$P_{Core3}$-DRE5-$P_{GAL1}$-yEGFP | This study |
| SAA3$^{C}$&2SCGy | EBY100 carrying pESD-AhR-ARNT-dCas9-$(SpyC)_2$-g3 and pBEVY-SpyT-Med2-*IGG6*-Gal3$^{C}$-$P_{Core3}$-DRE5-$P_{GAL1}$-yEGFP | This study |
| SAA3$^{C}$&3SCGy | EBY100 carrying pESD-AhR-ARNT-dCas9-$(SpyC)_3$-g3 and pBEVY-SpyT-Med2-*IGG6*-Gal3$^{C}$-$P_{Core3}$-DRE5-$P_{GAL1}$-yEGFP | This study |
| SAA3$^{C}$&4SCGy | EBY100 carrying pESD-AhR-ARNT-dCas9-$(SpyC)_4$-g3 and pBEVY-SpyT-Med2-*IGG6*-Gal3$^{C}$-$P_{Core3}$-DRE5-$P_{GAL1}$-yEGFP | This study |

**Table S4 Best fits for the characterized responses of the various sensors in this study**

| Sensors | Target | $Kc \times 10^{-7}$ (M) | $n$ | $k \times 10^{6}$ (a.u.) | LOD$\times 10^{-8}$ (M) | NF | $R^2$ |
|---|---|---|---|---|---|---|---|
| SAAC1y | β-NF | 3.56±0.58 | 0.89±0.078 | 0.74±0.027 | 4.59 | 1 | 0.9733 |
| SAAC13$^{C}$Gy | β-NF | 2.93±0.071 | 0.92±0.085 | 0.96±0.041 | 3.97 | 0.93 | 0.9820 |
| SAAC33$^{C}$Gy | β-NF | 2.51±0.16 | 1.73±0.040 | 2.06±0.19 | 3.24 | 0.18 | 0.9888 |
| SAAC43$^{C}$Gy | β-NF | 2.65±0.32 | 1.55±0.12 | 1.99±0.20 | 3.68 | 1.76 | 0.9911 |
| SAAC83$^{C}$Gy | β-NF | 2.85±0.20 | 1.31±0.10 | 1.54±0.15 | 3.34 | 1.79 | 0.9857 |
| SAA3$^{C}$&1SCGy | β-NF | 2.34±0.14 | 2.05±0.21 | 3.34±0.27 | 1.81 | 1.38 | 0.9948 |
| SAA3$^{C}$&2SCGy | β-NF | 2.37±0.21 | 1.95±0.086 | 2.67±0.18 | 2.03 | 1.19 | 0.9897 |
| SAA3$^{C}$&3SCGy | β-NF | 2.22±0.22 | 2.20±0.37 | 3.28±0.24 | 1.95 | 1.20 | 0.9962 |
| SAA3$^{C}$&4SCGy | β-NF | 2.16±0.12 | 2.34±0.34 | 3.75±0.20 | 1.05 | 0.85 | 0.9967 |

Data are presented as means, or means with 95% confidence intervals.

**Figure S1**

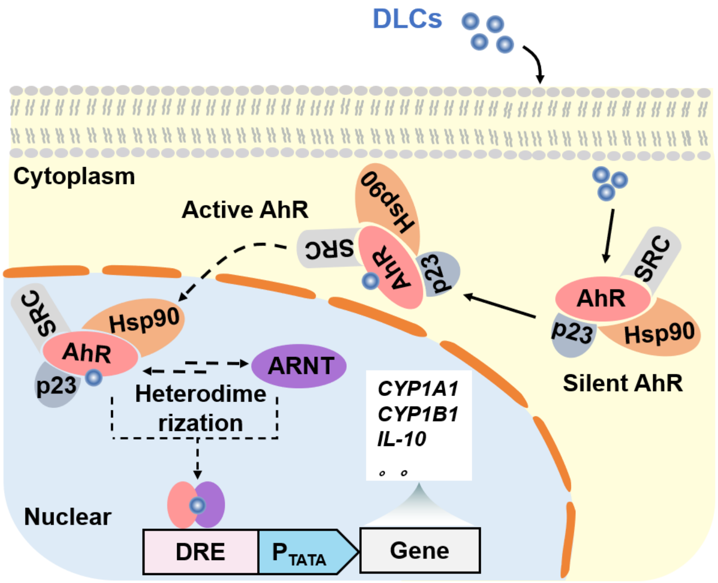


**Fig. S1** Schematic diagram of the AHR signaling pathway. The aryl hydrocarbon receptor (AHR), in its inactive state, complexes with the 90 kDa heat shock protein (HSP90), p23, and the protein SRC, among others, while maintaining it in a high-affinity conformation primed for ligand binding. Upon binding of dioxin-like compounds (DLCs), AHR undergoes exposes its N-terminal nuclear localization sequence (NLS) and subsequently heterodimerizes with its nuclear translocator ARNT to form a ternary DLCs–AHR–ARNT complex, which further binds to DNA-responsive elements (DRE) to control the expression of genes such as *CYP1A1*, *CYP1B1*, and *IL-10*.

**Figure S2**

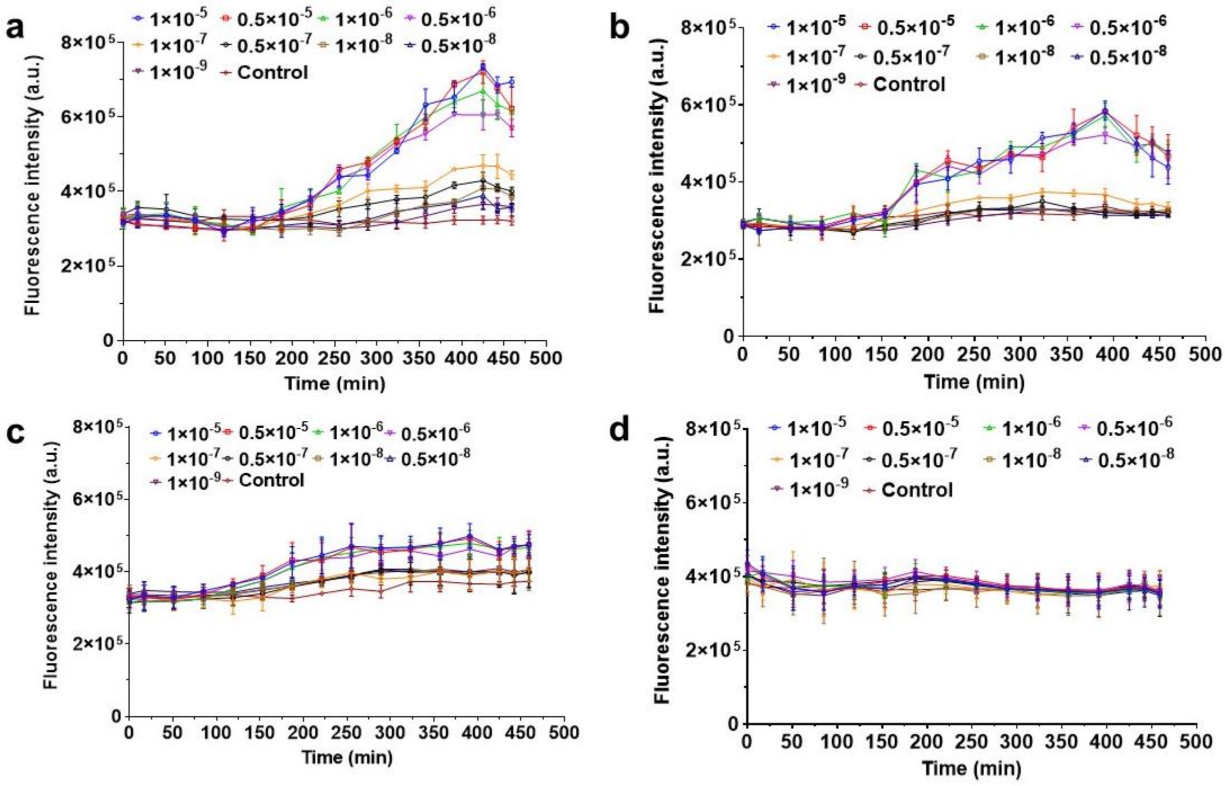


**Fig. S2** Time course of yEGFP signal output from engineered sensor strain **a**. SAAC1y, **b**. SAAC1y, **c**. SAAC3y, SAAC4y, and **d.** SAAC8y responding to different concentrations of β-NF. Data represent the mean and standard deviation of three independent experiments.

**Figure S3**

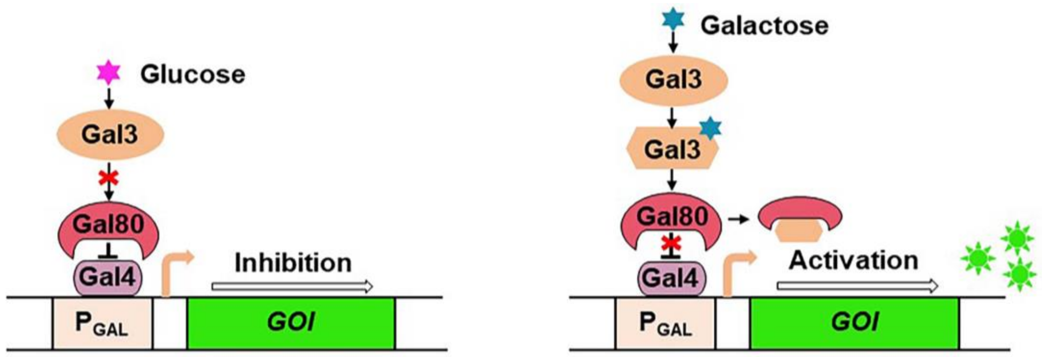


**Fig. S3** Schematic diagram of the *GAL* network. The transcriptional activator Gal4 binds to the promoter of the GAL gene. In the absence of galactose (glucose cultivation), the repressor Gal80 blocks the activation domain of Gal4, resulting in failure to initiate expression of downstream gene of interest (GOI). In the presence of galactose, Gal3 is activated, preventing Gal80 from binding to Gal4, resulting in transcription of GOI.

**Figure S4**

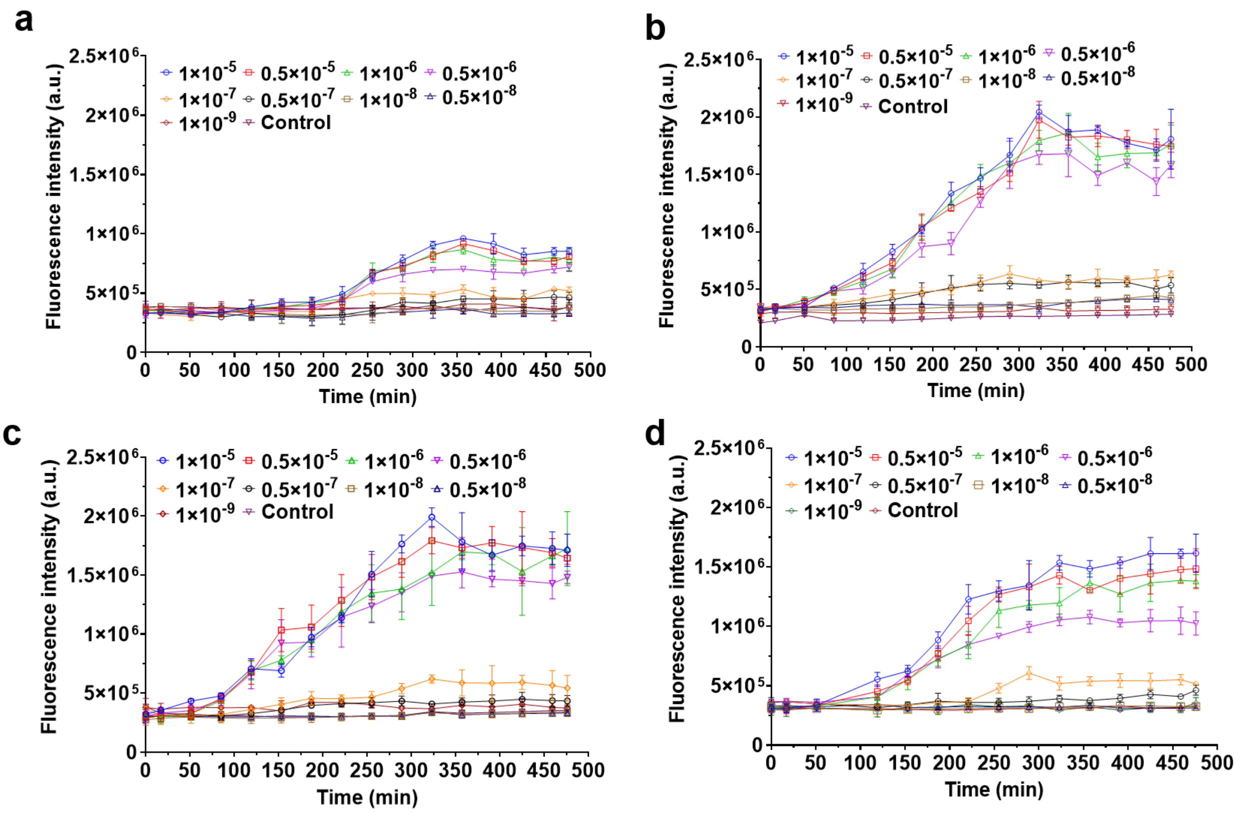


**Fig. S4** Time course of yEGFP signal output from engineered sensor strain **a**. SAAC13$^{C}$y, **b**. SAAC33$^{C}$y, **c.** SAAC43$^{C}$y, **d**. SAAC83$^{C}$y, responding to different concentrations of β-NF. Data represent the mean and standard deviation of three independent experiments.

**Figure S5**

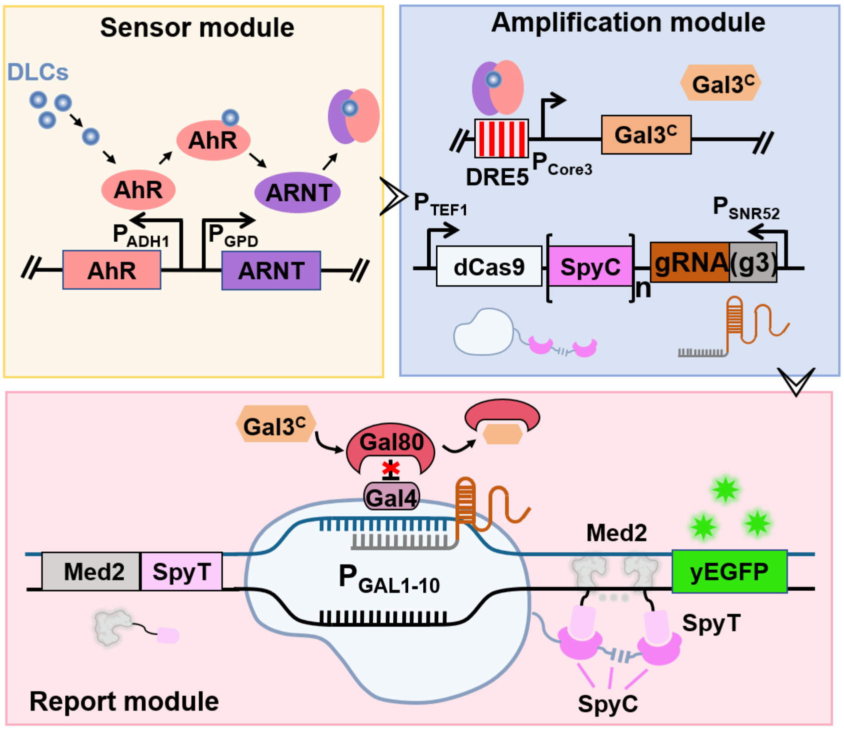


**Fig. S5** Schematic diagram showing the synergistic amplifier-based biosensor for DLCs response, including the sensor module (constitutive expression of AhR and ARNT), the amplification module (transcriptional activation component mediated by the Gal3$^{C}$ regulator and a transcriptional recruitment component mediated by dCas9-(SpyC)$_n$ complex), reporter module (simultaneous expression of yEGFP and TA-SpyT fusion). Once TA-SpyT is produced, it is captured by dCas9-(SpyC)$_n$ via SpyC-SpyT conjugation and targets the pre-defined DNA sequence of $P_{GAL1-10}$ to achieve higher expression of yEGFP.

**Figure S6**

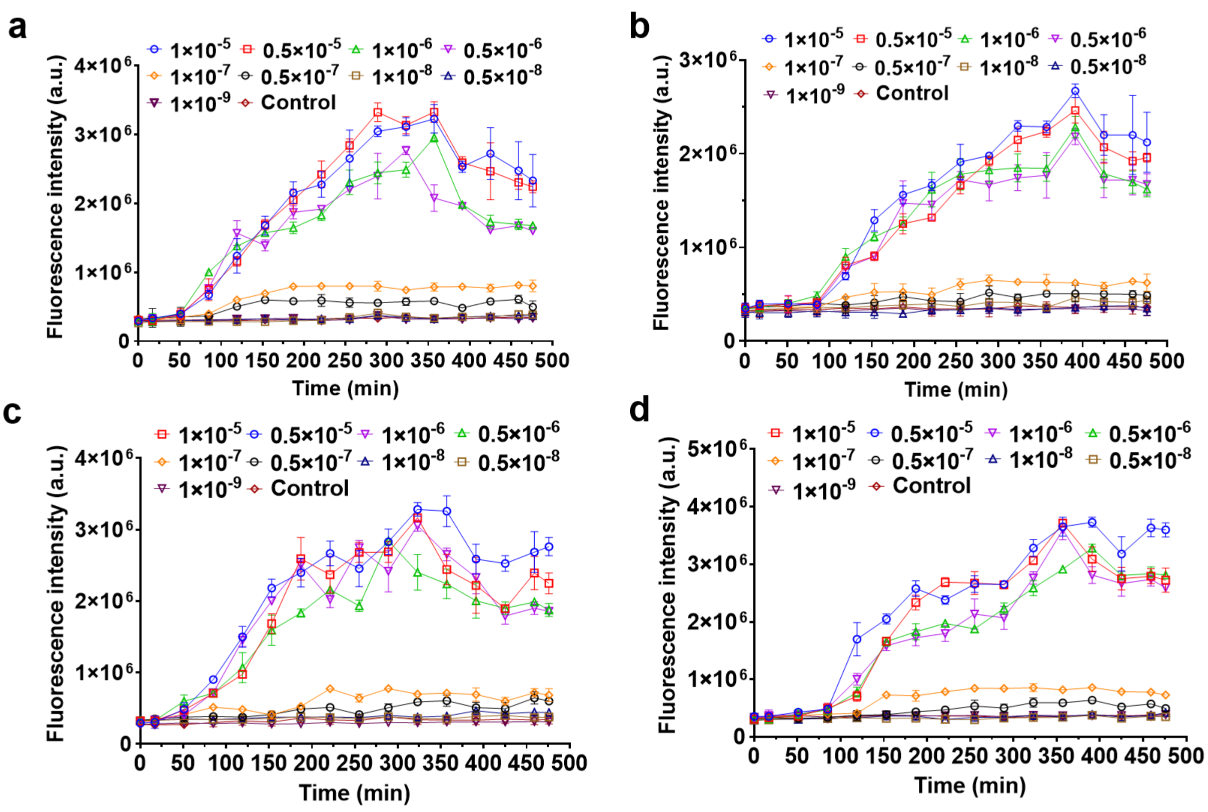


**Fig. S6** Time course of yEGFP signal output from engineered sensor strain **a**. SAA3$^{C}$&1SCGy, **b**. SAA3$^{C}$&2SCGy, **c**. SAA3$^{C}$&3SCGy, **d**. SAA3$^{C}$&4SCGy, responding to different concentrations of β-NF. Data represent the mean and standard deviation of three independent experiments.
